# Environmental surveillance of soil-transmitted helminths and other enteric pathogens in settings without networked wastewater infrastructure

**DOI:** 10.1101/2024.09.15.613066

**Authors:** Joël Edoux Eric Siko, Kendra Joy Dahmer, Zayina Zondervenni Manoharan, Ajithkumar Muthukumar, Heather K. Amato, Christopher LeBoa, Michael Harris, Venkateshprabhu Janagaraj, Malathi Manuel, Tintu Varghese, Parfait Houngbegnon, Nils Pilotte, Bernadin Bouko, Souad Saïdou, Adrian J. F. Luty, Rohan Michael Ramesh, Moudachirou Ibikounlé, Sitara S.R. Ajjampur, Amy J. Pickering

**Author notes:** equal contributors as co-first authors (Joël Edoux Eric Siko, Kendra Joy Dahmer).

## Abstract

Soil-transmitted helminths (STH) are one of the most prevalent enteric infections world-wide. To control STH-related morbidity, the World Health Organization recommends targeted deworming and improvements in water, sanitation and hygiene. Current surveillance strategies for STH focus on identifying and quantifying eggs in stool samples via microscopy, which exhibits poor specificity and sensitivity, especially in settings with low-intensity infections. Wastewater-based epidemiology is a surveillance tool used to monitor pathogen circulation and could replace stool-based approaches for STH detection. However, sampling strategies for settings lacking networked sanitation outside large urban settlements are not well developed. Here, we report evaluation of sampling strategies for soil and wastewater STH surveillance in rural and peri-urban settings without networked sanitation. We used multi-parallel qPCR assays to detect STH DNA in soil collected from high foot-traffic locations and three types of wastewater samples (passive Moore swabs, grab samples, and sediment from drainage ditches) in Comé, Benin and Timiri and Jawadhu Hills in Tamil Nadu, India. We detected STH in soil (India = 32/95, Benin = 39/121) and wastewater (India = 24/60, Benin = 8/64) with a detection frequency across all sample types of 36% in India and 25% in Benin. We evaluated which sample locations and types allowed for more sensitive detection of STH DNA and determined that STH prevalence varied by sample site but did not vary significantly within a given sample site location (e.g., samples collected from multiple locations within one market). Further, we determined that wastewater sediment samples outperformed grab and Moore swab sample types for STH detection. Finally, we expanded our methods to include detection of other enteric pathogens using multiplexed qPCR for wastewater samples. Our results establish sampling strategies for environmental and wastewater surveillance of a wide range of enteric pathogens in settings without networked sanitation.

## 1 Introduction

Soil-transmitted helminths (STH) account for over five million disability-adjusted life years (DALYs) and infect around 1.5 billion people, representing the most prevalent parasitic infections worldwide (1). *Ascaris lumbricoides*, *Trichuris trichiura*, *Necator americanus,* and *Ancylostoma duodenale* are the most common human STH, but other species are known to have zoonotic potential (2). Infections with STH cause several health problems, such as malnutrition, anemia, poor birth outcome (low birth weight, mortality), and child developmental deficits (3–7). STH infection and transmission occur primarily in tropical and subtropical regions and are inextricably linked to poverty, inadequate sanitation, and unhygienic conditions.

Transmission of STH occurs when eggs, that are passed in human feces, contaminate the soil, leading to onward infections through the ingestion of eggs or via hookworm infective larvae (iL3’s) penetrating the skin. Current STH control and elimination (as a public health problem) programs rely on mass drug administration (MDA) with monitoring of intervention success conducted using human stool-based methods (Kato-Katz microscopy) (8). MDA campaigns for STH mainly focus on treating school-aged children in endemic countries and have had varied success with reinfection frequently and rapidly occurring (7,9,10). Microscopy-based estimates of STH prevalence are limited by day-to-day variation in helminth egg excretion, parasite egg misidentification, and compliance of stool donors (11). Obtaining stool samples is often difficult, particularly from adolescent and adult populations, due to stigma, reluctance, and lack of perceived benefit (12). Elimination or sustained transmission interruption of STH has not occurred through MDA alone and elimination typically requires improved infrastructure likely due to the environmental life-cycle requirements and potential environmental reservoirs of infectious stage STH (13–16).

Wastewater-based methods are now widely used for surveillance of pathogen circulation and to identify potential reservoirs of disease transmission. The recent COVID-19 pandemic has shown the utility of such approaches using sewage samples to track disease prevalence (17,18). Translating the success of wastewater-based epidemiology to environmental surveillance tools for STH may provide a non-invasive, more cost-effective solution to environmental monitoring of these pathogens. However, wastewater surveillance typically relies on the networked sanitation systems found in high-income settings, where wastewater treatment plants can provide samples that are representative of a population based on its catchment area. Less is known about the feasibility and utility of wastewater surveillance of pathogens in low- and middle-income countries (LMIC), where sewage is not systematically treated and may be contained in pits or septic tanks, discharged in storm drains, or discharged directly into the environment.

STH are endemic to both India and Benin, with around 20% overall population prevalence and with hookworm infections predominant (19,20). Since it remains unclear where hotspots for transmission and environmental reservoirs of STH are located in these endemic communities (21), we sought to determine the utility of public soil and wastewater environmental DNA (eDNA) sampling as a surveillance tool for STH in communities in both India and Benin that lack networked wastewater sanitation infrastructure. Using molecular surveillance tools to identify STH has been shown to be more species-specific (differentiating between human and animal parasite eggs or larvae) and more sensitive than Kato-Katz microscopy (22). qPCR has emerged as the best molecular tool for identifying polyparasitism since it is easily and cheaply multiplexed or parallelized and gives highly sensitive quantitative readouts. Here, we use multi-parallel qPCR assays to detect STH eDNA from a variety of soil and wastewater sample sites and multiplexed qPCR to detect additional enteric pathogens of clinical importance in wastewater from sites in Comè, Benin and Tamil Nadu, India with a focus on methods of acquiring environmental samples and validating detection of eDNA in communities lacking networked sanitation infrastructure.

## 2. Materials and Methods

### 2.1 Field Collection of Soil and Wastewater Samples

Soil and wastewater samples (Benin: soil n = 121, wastewater = 64, India: soil n = 95, wastewater n = 60) were collected from five previously defined control clusters (23,24) enrolled in an ongoing cluster randomized trial in each country where school-aged children received annual (Benin) or biannual (India) deworming medication (Albendazole) as per the standard of care. In each cluster, soil from two markets (three samples each: entrance, center, path), two schools (three samples each: entrance, classroom, path to latrine), two open defecation fields (three samples each: entrance, center, field edge), and one (India) or two (Benin) community water point(s) were collected. Additionally, soil samples were collected from five households per cluster in Benin. Three types of wastewater samples (a grab, a sediment, and a Moore swab) from two wastewater drainage sites per cluster, in each country, were also collected. Field staff identified sampling locations using a password protected tablet or smartphone loaded with the SurveyCTO form containing GPS coordinates for the relevant country and cluster.

#### 2.1.1 Soil Collection

For soil sampling, methods were adapted from previous studies conducted by our group (25). Briefly, a 30 cm x 50 cm disposable soil stencil was laid out on the sampling area and the top surface soil equating to roughly 100 grams from inside the stencil area was scraped using a sterile scoop and scooped into a Whirlpak bag (WPB01350WA, Merck). The bag was sealed by folding along the wire strips and flaps then labeled with the corresponding label type after wiping down the bag with 70% ethanol and 10% bleach. The Whirlpak bags with soil samples were stored in a cooler box and the scoop and stencils were discarded. The soil samples were sent to the lab in the cooler box and kept at 4°C until processing. Samples were processed within 24 hours of being received in the laboratory.

#### 2.1.2 Wastewater Collection

For wastewater grab sampling, a sterile 500 mL Whirlpak bag (WPB01350WA, Merck) was slowly immersed into the flowing wastewater channel allowing the bag to fill with wastewater. For wastewater sediment sampling, one field staff member held a sterile Whirlpak bag while another field staff member scraped 250 mL of wet sediment from the bottom of the channel with a sterile scoop. For Moore swab samples, a 4x4 ply gauze (7086, ExcilonTM) was tied with a fishing line (00291073, Shur Strike). The fishing line anchored the swab in the channel and the other end was tied to a branch or floating object such that the swab was floating in the middle of the water channel to allow for wastewater filtering. The Moore swab was left suspended in the stream of wastewater for 24 hours before the string was cut and the swab was placed into a sterile Whirlpak bag (WPB01065WA, Merck). All bags were sealed by folding along the wire strips and flaps. Bags were then wiped down with 70% ethanol and 10% bleach, after which they were appropriately labeled. At the end of each day, one field blank sample was prepared whereby a sterile Whirlpak bag was filled with clean bottled water in the field and labeled as ‘blank’. All samples were sent to the lab in a cooler box and held at 4°C until processing. Samples were processed within 24 hours of being received in the laboratory.

### 2.2 Sample Laboratory Processing and Analysis

Upon receipt from the field, samples were scanned in the laboratory using the appropriate SurveyCTO form and processed data was entered into the appropriate processing SurveyCTO form.

#### 2.2.1 Soil Processing

For each soil sample, soil was sieved through a 2 mm mesh screen to get rid of rocks and other debris. The sieved soil was mixed thoroughly and 40 gram aliquots were scooped into 50 mL centrifuge tubes. The 50 mL centrifuge tubes with sieved soil samples were labeled with barcodes, parafilmed (1337412, Fischer) around the cap, and stored at -80°C until DNA extraction. Reusable soil screens were cleaned before reuse by first washing with soap followed by soaking in 70% ethanol for 2 minutes and air drying.

#### 2.2.2 Wastewater Processing

Wastewater grab samples were vacuum filtered (EZFITMIHE1, Merck) through a 0.45 µm filter paper (MIHAWG250, Merck). Whirlpak bags were gently shaken before 250 mL of liquid was transferred to the filter. After 30 minutes of filtering, if the liquid had not fully passed through, the remaining liquid was aspirated and transferred to a new filter paper, repeating as needed for up to four filter papers per sample. After the entire sample (500 mL) had been filtered or before switching to a new filter, the manifold was closed and 500 µL of Zymo RNA/DNA Shield (R1100250, Zymo) was pipetted over the entire surface of the filter and incubated for 30 seconds. The vacuum was opened and the RNA/DNA shield was allowed to flow through. Using sterile forceps, the filter paper was carefully removed, rolled up and placed in a 5 mL tube. If there was more than one filter paper per sample, all filters were stacked on top of each other in a petri-dish, rolled together and placed in the same 5 mL tube. The tube was parafilmed, labeled appropriately, and placed at -80°C until DNA extraction.

For processing wastewater sediment samples a small hole was cut in the bottom corner of the Whirlpak bag with sterile scissors and the liquid was allowed to drain out. 45 mL of wet sediment was then transferred into a sterile 50 mL tube. The tube was labeled with a barcode, wrapped with parafilm, and stored at -80°C until DNA extraction.

For processing wastewater Moore swab samples, sterile forceps were used to remove the Moore swab from the Whirlpak bag. Using sterile scissors the swab was cut in half and each half was placed into separate sterile 50 mL tubes. 30 mL of 1X PBS (20012043, ThermoFisher) was added to both tubes. Tube one was vortexed for 10 minutes at maximum speed. The swab was then removed using ethanol flame sterilized forceps. The tube was centrifuged for five minutes at 1000 x g and the supernatant was discarded. The pellet at the bottom of the tube was resuspended in 1 mL of nuclease free water and transferred to a 2 mL storage tube. The 2 mL tube was labeled with a barcode, parafilmed, and stored at -80°C until DNA was extracted using the DNeasy PowerSoil Pro kit (47014, Qiagen). Tube two, containing half of the Moore swab in 30 mL of 1X PBS, was labeled and stored at -80°C until DNA was extracted using the DNeasy PowerMax Soil kit (12988-10, Qiagen).

### 2.3 DNA Extractions

All soil samples and the wastewater sediment samples were processed using the DNeasy PowerMax Soil kit (Qiagen cat # 12988-10). The DNeasy PowerWater kit (Qiagen cat # 14900- 100-NF) was used to process wastewater grab samples. Moore swabs were cut into two pieces and processed using either the DNeasy PowerSoil Pro Kit (Qiagen cat # 47014) (Tube 1) or the DNeasy PowerMax Soil Kit (Qiagen cat # 12988-10) (Tube 2). An extraction blank (reagent only extraction), negative control for ensuring no cross contamination occurred, was performed for every 35 soil samples in India and every 30 soil samples in Benin for soil samples that were processed.

#### 2.3.1 Soil, Wastewater sediment and Moore swab (direct extraction)

DNA extractions of soil, wastewater sediment, and one half of the Moore swab samples was performed using the DNeasy PowerMax Soil Kit (1298810, Qiagen) in accordance with the manufacturer’s protocol with some modifications (25). Briefly, input soil quantity was 20 grams per extraction. For Moore swab tube number two (T2), stored in 30 mL of 1X PBS, we added 15 grams of powerbeads to the tube and vortexed for 10 minutes. The swab was then removed and DNA extraction proceeded on the eluant. Modification to the extraction protocol included: samples were vortexed for 30 minutes at maximum speed (Vortex Genie 2, Scientific Industries, Cat# 2E- 242275) in the PowerMax Bead Pro Tube followed by only half of the supernatant being transferred to a clean collection tube. After adding the solution C4, the internal control was added to the tube. Internal controls were 5 µl of *Bacillus atrophaeus* (0801824, ZeptoMetrix LLC) diluted at 1:100 (initially titered at 1.30 x 10^9^ CFU/ml) in Benin and 1 µl of Internal Amplification Control (IAC), pDMD801 (26), a synthetic and non-coding in-house plasmid, at 100 pg/µl in India. Final DNA elutions were concentrated and precipitated immediately after extraction.

To concentrate and precipitate the DNA elution, 5 µL of Pellet Paint NF Co-precipitant (Sigma Millipore Cat. # 70748-3) and 500 µL of Sodium Acetate (pH 5.2) was added to each sample elute. 10 mL of 100% ethanol was added to each sample, vortexed briefly and incubated at room temperature for two minutes. The tube was centrifuged at top speed for five minutes. Without disturbing the pellet, the supernatant was discarded and 10 mL of 70% ethanol was added to the pellet. The pellet was vortexed and centrifuged at top speed for five minutes. The supernatant was again discarded and 10 mL of 100% ethanol was added to the pellet. The pellet was vortexed and centrifuged at top speed for five minutes. The supernatant was removed and the pellet was dried overnight in a biosafety cabinet or on a heat block at 90°C for five to ten minutes per sample. After drying the pellet, 100 µL of nuclease-free water was added to resuspend the pellet. After resuspension, the entire 100 µL of concentrated DNA was transferred to a clean cryovial and stored at -80°C.

#### 2.3.2 Wastewater Grab

The DNeasy PowerWater kit (1490050-NF, Qiagen) was used for extraction of DNA from wastewater grab samples as per the manufacturer’s protocol with the following modifications: The filters from grab sample processing were transferred to the 5 mL PowerWater DNA bead tube provided. After adding the solution PW3, the internal control was added to the tube. Internal controls were the same as those used with the Qiagen Dneasy PowerMax Soil Kit as described above. Eluted DNA was stored at -80°C.

#### 2.3.3 Moore Swab Pellet

The pellet from the first Moore swab tube (T1) was extracted using the DNeasy PowerSoil Pro Kit (12988-10, Qiagen) following the manufacturer’s protocol with the following modifications: After adding the solution CD3, the internal control was added to the tube. Internal controls were the same as those used with the Qiagen DNeasy PowerMax Soil Kit as described above. Final eluted DNA was stored at -80°C.

### 2.5 qPCR Procedure for multi-parallel Real-time PCR Analysis

qPCR was performed in duplicate for the identification of STH eDNA using species-specific primers and probes targeting repetitive sequences (25,27,28) (**S1 table**). Samples were considered positive if at least one of the replicates produced a PCR product with a Cq value <40. We removed samples with failed internal amplification controls (IACs) from our analysis **(S1, S2 and S3 Fig).**

#### 2.5.1 qPCR for Samples Collected in India

In India, qPCR was performed as previously described (25). Briefly, multi-parallel qPCR assays targeting non-coding repetitive sequences were utilized to detect *Necator americanus*, *Trichuris trichiura*, and *Ascaris lumbricoides* (**S1 Table**) in both soil and wastewater samples. All samples were tested in duplicate and a standard dilution series of plasmid controls (10 pg, 100 fg, and 1 fg) containing a single copy of the target sequence for each assay was utilized as a positive PCR control. No template control (NTC) samples were tested on each qPCR reaction plate. Each sample was also tested for the IAC (pDMD801) (26) to ensure that DNA recovery occurred during extraction and that recovered DNA was amplifiable (**S1 and S2 Fig**). Individual aliquots of TaqPath ProAmp mastermix (A30872, ThermoScientific) and species-specific primers and probe (Integrated DNA Technologies) stocks were prepared before setting up the qPCR assay. Mastermix for a plate of 96 reactions was assembled by adding 350 µL of TaqPath ProAmp mastermix (A30872, ThermoScientific), primers and probe at 10 µM concentrations, and nuclease free water to a final total volume of 530 µL. 5 µL of the prepared mastermix was then aliquoted into each well of the 96-well plate (4306737, ThermoFisher). 2 µL of the DNA template (environmental/wastewater sample eDNA or plasmid dilution) was then added to each well.

For all the STH assays, cycling conditions included an initial two minute incubation step at 50°C, followed by a ten minute incubation at 95°C, then 40 cycles of: 15 seconds at 95°C for denaturation and one minute at 59°C for annealing and extension. All qPCR reactions were carried out using the Quantstudio 7 Flex PCR system (Applied Biosystems) and the data generated was analyzed using QuantstudioTM Real-Time PCR software Version 1.3. A sample with a Cq value <40 in one or both replicates was reported as positive for the target tested. Field blank samples were included as negative controls (**S3 Fig**).

#### 2.5.2 qPCR for Samples Collected in Benin

In Benin, lyophilised plates (221115017-221117006, Argonaut) containing one set of species-specific primers, probe, and mastermix for *N. americanus*, *A. duodenale*, *T. trichiura*, or *A. lumbricoides* were used for qPCR (**S1 Table**). Each DNA sample was run in duplicate. Each reaction plate contained four controls including an IAC (*B. atrophaeus* added during extraction and a known positive sample for *A. lumbricoides*), an extraction negative control (no sample added during extraction), a qPCR negative control (NTC), and a qPCR positive standard control. The qPCR positive control was a single plasmid (1 fg) containing the target sequences for each target species (**S1 and S2 Fig**). Prior to running the lyoplates, each DNA sample was diluted by adding 6 µL of molecular grade water to each well of a dilution plate followed by adding 18 µL of DNA sample to each well. Resuspension of lyocakes inside each well of the lyoplate occurred by adding 10 µL of diluted DNA sample. All qPCR assays were performed using the QuantStudio 5 real-time PCR system (Applied Biosystems). Cycling conditions were an initial two minute incubation step at 50°C, followed by a ten minute DNA denaturation at 95°C, then 40 cycles of: 15 seconds at 95°C for denaturation and one minute at 59°C for annealing and extension. A sample with a Cq value <40 in one or both replicates was reported as positive for the target tested. Field blank samples were included as negative controls (**S3 Fig**).

### 2.6 Real-time PCR using TaqMan Array Cards (TAC)

The TaqMan array card (TAC) assay was used for the detection of multiple enteropathogen targets in wastewater samples (**S3 Table**). The TAC assay was a 384-well (Benin) or 48-well (India) (29,30) arrayed singleplex real-time PCR assay which was used for the simultaneous detection of 30 enteric pathogens in Benin and 18 enteric pathogens in India. The enteric TAC targets include bacteria, viruses, protists and helminths and the universal bacterial 16S target for normalization of pathogen burden to total bacterial load. 40 µL of DNA from each sample was mixed with 60 µL of the mastermix comprised of AgPath-ID One step RT PCR buffer, enzyme mix (containing reverse transcriptase and Taq polymerase enzymes) and nuclease free water (4387391, ThermoFisher). The 100 µL reaction mixture was loaded onto each sample port of the TAC card and centrifuged at 4°C for 2 minutes at 1200 rpm. The plate was sealed with an optical adhesive sealer (4311971, ThermoFisher) and the sample ports were excised. The PCR was assayed on the QuantStudio 7 flex with the pre-set run template and the cycling condition described in **Table 2**. Analysis was carried out using the QuantStudio 7 software.

### 2.7 Data availability and Statistical analysis

All qPCR data and pipelines for statistical analysis and data visualization were performed using R statistical software (version 4.3.2) and are available at https://github.com/KendraDahmer/wastewater-ms. To compare detection prevalence between soil and wastewater samples, we performed Fisher’s exact test with odds ratio (OR) and 95% confidence intervals (CI). To distinguish between specific sample types and sites and compare species-specific composition, we performed Fisher’s exact pairwise tests (**S1 data**).

## 3 Results

### 3.1 Sampling strategy

We collected, processed and analyzed 185 samples from Benin in the community of Comé and 155 samples from Tamil nadu in India (Timiri and Jawadhu Hills) (**Fig 1 A and B**). Soil samples were collected from markets, schools, households, open defecation fields, and community water points. Wastewater samples were collected from open storm drains with flowing wastewater when available, or from stagnant wastewater in drains or at points of discharge. Soil samples were collected within a 30 cm x 50 cm stencil by scraping once vertically and then once horizontally throughout the stencil to acquire 100 grams of topsoil (**Fig 1 C**). At each wastewater sampling location three sample types were collected including a 300 mL water grab sample, a 250 gram sediment sample scraped from the bottom of the waterway, and one Moore swab (**Fig 1 D**). Four- ply Moore swabs were suspended in the middle of the water column for 24 hours in order to acquire composite samples. To assess species presence and which sample type would likely be most valuable for environmental surveillance, we used species-specific qPCR assays.

**Fig 1.**
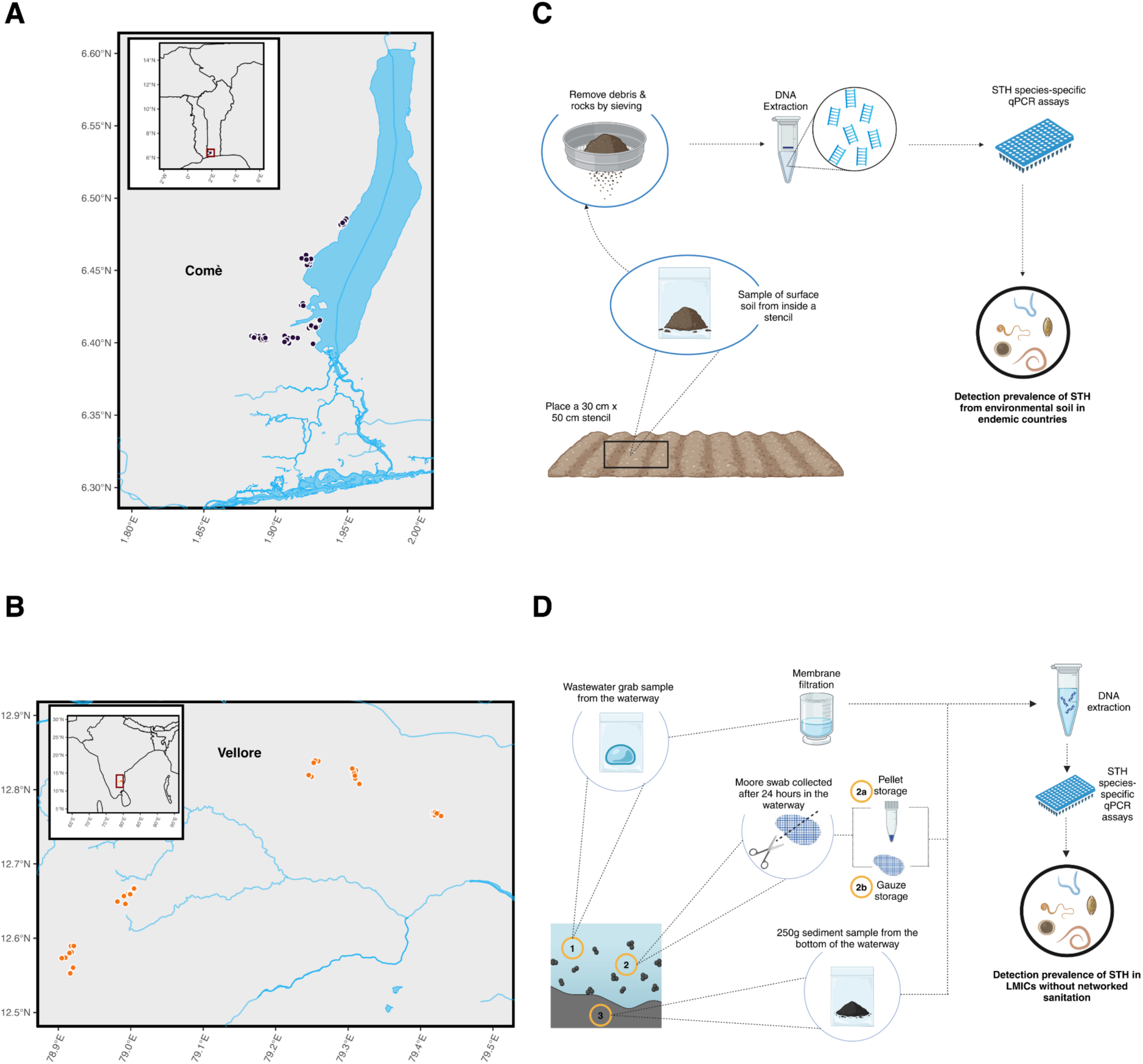
**Sample collection sites and sampling methods**. A-B) Map of sampling sites in Benin and India, respectively. Five clusters were sampled in each country for detection of STH. C) Methods overview of soil sample collection for STH detection using qPCR. Top soil was scraped from within a 30 cm x 50 cm stencil and 100 grams of soil was collected for DNA extraction and qPCR. D) Methods overview of wastewater sample collection in countries without networked sanitation systems for STH detection using qPCR. Three wastewater sample types were collected from each sampling location. Moore swabs were cut in half and processed with two different methods (2a and 2b) (see methods for details).

### 3.2 STH detection in soil and wastewater samples

We first sought to validate detection of STH within our study sites of Comé, Benin and Timiri and Jawadhu Hills, Tamil Nadu, India. For samples collected in Benin, qPCR was performed to identify the presence of *A. lumbricoides*, *T. trichiura*, *N. americanus,* and *A. duodenale* eDNA in environmental samples. For samples collected in India, qPCR was performed to identify the presence of *A. lumbricoides*, *T. trichiura*, and *N. americanus* eDNA in environmental samples. We identified 47 samples (25%) positive for STH in Benin and 56 samples (36%) positive for STH in India (**Fig 2 A**). *T. trichiura* was not detected in any sample in either country. *N. americanus* had the highest prevalence (29.03%) followed by *A. lumbricoides* (10.3%) in environmental samples from India, while *A. lumbricoides* (14.05%) was the most prevalent in Benin (**Fig 2 B**). Consistent with stool sampling data from previous studies in our study areas (19,31), hookworm (*N. americanus* and *A. duodenale* combined) was the most prevalent eDNA identified in our soil and wastewater samples.

**Fig 2.**
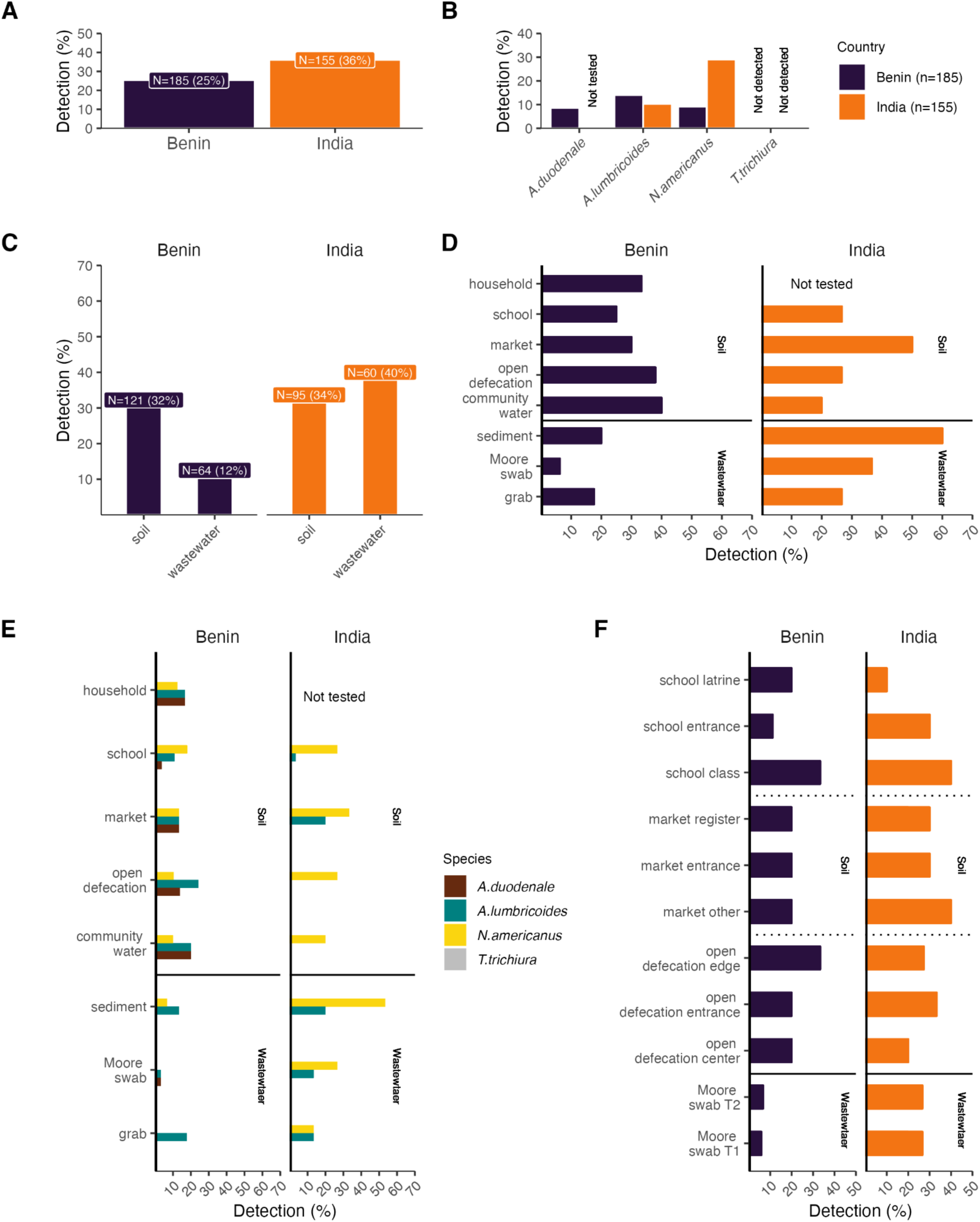
Prevalence of STH species by study site and sample type as determined by qPCR. A) Overall detection prevalence of STH in each of our study countries. B) Prevalence of STH species in each study country. C) Prevalence of any STH species in soil and wastewater sample types. D) Prevalence of STH in each sampling site. E) Breakdown of prevalence of STH species in each sample site. F) Prevalence of any STH species in each sampling site and location and prevalence in Moore swabs processed with two different methods. T1 = tube one, T2 = tube two. Fisher’s exact test p-values can be found in **S1 data.** Benin = purple, India = orange throughout the figure. No *T. trichiura* was detected in any samples from India or Benin.

In order to optimize a reliable sampling strategy, we sought to evaluate which sample types allowed for more sensitive detection of STH eDNA. We determined that our overall ability to detect any STH species was not significantly different between soil and wastewater samples collected in both countries combined (p-value = 0.18, OR = 1.4064, 95% CI = 0.8393, 2.3886). However, detection differed significantly between soil and wastewater samples collected in Benin (p-value = 0. 0042, OR = 3.310, 95% CI = 1.38904, 8.8308), with soil samples having higher detection prevalence, but not in India (p-value = 0.4931, OR = 0.7633, 95% CI = 0.3707, 1.5761) (**Fig 2 C**). Further, we determined that Moore swabs from Benin were significantly less likely to detect STH than most soil samples (p-values: household = 0.0133, markets = 0.02, open defecation = 0.00393, community water = 0.0214) (**Fig 2 D**). In India, soil samples from open defecation fields (p-value = 0.0497) and schools (p-value = 0.0497) were significantly less likely to detect STH compared to wastewater sediment samples. To further evaluate and optimize a sampling strategy that would provide a robust view of species-specific prevalence at each sample collection site, we evaluated whether species-specific detection was significantly different between sample types and if sampling location within a site altered our ability to detect STH species. We determined that species-specific composition differed significantly between some sample sites (**Fig 2 E**). *N. americanus* was undetected in wastewater Moore swabs in Benin, while detection occurred from soil samples collected at schools (p-value = 0.018) and markets (p-value = 0.0491). *A. lumbricoides* was detected more frequently in soil from open defecation fields than in wastewater Moore swabs (p-value = 0.022). In India, *A. lumbricoides* was undetected in soil samples collected from open defecation fields while detection occurred from wastewater sediment samples (p-values = 0.0321) and soil samples collected from markets (p-value = 0.0237).

In both countries, detection of any STH species did not significantly differ between the within site locations (**Fig 2 F**). Furthermore, processing differences for Moore swabs did not result in significant differences in STH detection. Overall, there was agreement between STH detection rates across sample sites and locations and within sample sites.

### 3.3 TaqMan Array Cards for diverse enteric pathogen detection

In order to expand our surveillance efforts to include bacterial, viral and single-celled parasitic enteric pathogens in addition to STH and to optimize methods for wastewater surveillance in LMICs lacking networked sanitation, we used TaqMan Array Cards (TAC) to simultaneously detect a wide range of enteric pathogens. eDNA from wastewater grab samples (India n = 15, Benin n = 2), sediment samples (India n = 4, Benin n = 8), and Moore swabs (India n = 15, Benin n = 6) as well as a subset of soil samples from Benin (n = 4) were assayed for the presence of a variety of enteric pathogens (enteric pathogens: India n = 18, Benin n = 30) (**S3 Table**). In Benin, TAC assays on soil samples only detected 4/30 enteric pathogens, while assays on wastewater sediment samples detected 13/30 enteric pathogens, wastewater Moore swab samples detected 12/30 enteric pathogen and wastewater grab samples detected 16/30 enteric pathogens in at least one sample tested of each type (**Fig 3 A**). Further, the type of pathogen detected varied across wastewater samples. Wastewater sediment samples detected more parasitic pathogens (4/8 parasites detected) while wastewater grab samples detected more bacterial pathogens (12/17 bacteria detected) than other sample types assayed. In India, wastewater sediment and Moore swabs detected more parasitic pathogens (4/4 parasites detected) than grab samples. Wastewater samples outperformed soil across all pathogen types. In India, TAC assays on wastewater sediment samples resulted in the detection of 11/18 enteric pathogens, wastewater Moore swab samples detected 16/18 enteric pathogens, and wastewater grab samples detected 14/18 enteric pathogens in at least one sample tested of each type (**Fig 3 B**). Wastewater Moore swabs and grab samples from India detected more viral pathogens (4/5 viruses detected) and bacterial pathogens (8/9 bacteria detected) than wastewater sediment samples. Overall, combining soil and wastewater sample collection lends an advantage to diversify enteric pathogen surveillance in LMICs lacking networked sanitation infrastructure (**Fig 3 C**).

**Fig 3.**
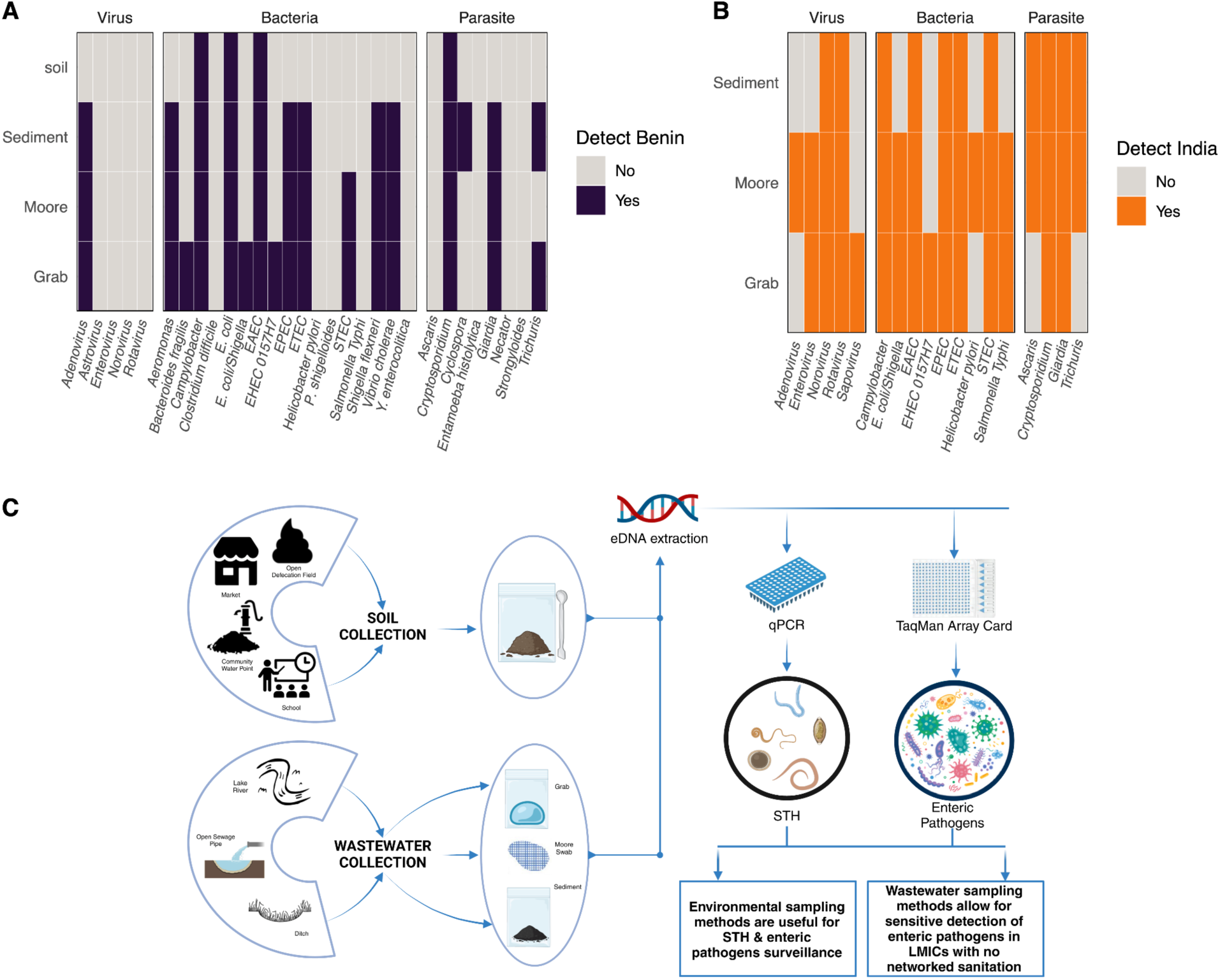
**Detection of enteric pathogens from non-networked wastewater using TaqMan Array Cards and optimized pipeline for sample collection and molecular surveillance**. A) Presence or absence of enteric pathogens (bacteria, viruses, parasites) in at least one sample of each sample type collected in Benin. B) Presence or absence of enteric pathogens (bacteria, viruses, parasites) in at least one sample of wastewater sample types collected in India. C) Schematic showing optimized steps for sample collection from soil and wastewater samples and an overview of processing and analysis steps for enteric pathogen detection. Benin = purple, India = orange

## 4. Discussion

We used molecular methods to detect STH and a prioritized list of clinically relevant enteric pathogens from soil and non-networked wastewater samples. Overall, we detected STH in 25% of samples from Benin and in 36% of samples from India. Soil collected from public locations with high foot traffic had similar prevalence of STH as wastewater samples collected from the same communities in India, and had higher prevalence than wastewater samples in Benin. These results suggest that sampling both soil and wastewater may be the most comprehensive approach to monitoring STH with environmental sampling, particularly when wastewater sites are seasonal or heterogeneous across communities.

One benefit of environmental sampling appears to be the widespread distribution of STH eDNA when present in a community sampling site (e.g. school, market). We determined there was no significant difference in detection of STH within site locations (e.g., school entrances, school classrooms or school latrine entrances) likely due to the widespread distribution of STH throughout soil from sites that may harbor environmental reservoirs of infectious stage STH (eggs, iL3s). These results suggest that collecting soil at a single location within a site is sufficient for surveillance efforts.

Despite having small sample sizes for some soil and wastewater sample types, molecular detection of each species was in relative agreement with recent trends of human infections based on Kato-Katz microscopy in Benin and India (19,20). qPCR analysis often results in higher detection frequency than Kato-Katz microscopy due to the high sensitivity and specificity associated with molecular detection (22,32,33). Given our data depicts a higher prevalence in the environment compared to human prevalence, environmental sampling could be a sensitive method for monitoring community-level infection prevalence. However, paired environmental and human stool sample collection is needed to determine how well environmental sampling can predict human infection prevalence at the community level.

In this study, wastewater sample types (Moore swabs, grab samples, and sediment) performed differently in Benin and India but will likely have utility in diverse LMICs depending on the type of infrastructure in place. Moore swabs are a reliable passive sampling tool optimized for use in flowing wastewater (34). However, stagnant wastewater sites are common and may not be appropriate for Moore swab capture, as the settling velocity of STH eggs can be on the order of 1 cm/minute (35). *Ascaris* eggs settle quicker than hookworm eggs and are more hardy (35), which affects their persistence and transport in the environment. In Benin, Moore swabs had lower detection of STH compared to soil samples and other wastewater samples. One explanation could be that our wastewater sites in Benin were primarily stagnant runoffs from a small number of households or runoff flowing into larger bodies of water such as a lake. Study communities in Benin lacked wastewater infrastructure, with wastewater draining through small pipes or ditches depositing into a puddle, a carved out crevice in the soil, or cement troughs. In our India study sites, wastewater infrastructure included cemented wastewater channels with stagnant or flowing water, roughly dug channels with running wastewater, or stagnant wastewater collections from households. Sediment samples were the most sensitive wastewater sample type for STH across both countries for the detection of STH, suggesting sediment should be prioritized for incorporation into STH environmental surveillance efforts.

Wastewater samples also allow for more diverse surveillance of enteric pathogens including bacteria, viruses and protists (36). Different wastewater samples were better at detecting specific groups of pathogens likely due to settling rate of the pathogen and factors such as quality of infrastructure, water depth and flow rate, and climate variables (34). Compared to soil in Benin, wastewater samples were able to capture a broader set of enteric pathogens, including bacterial, viral, and parasitic targets despite these samples originating from non-networked wastewater systems. Grab samples and Moore swabs were better able to capture bacteria and viruses compared to wastewater sediment samples.

For children in LMICs, diarrheal disease is among the leading causes of mortality (37) while STH infections are among the leading causes of chronic morbidity (7,38,39). While soil and sediment were the most sensitive approach for an STH focused environmental surveillance strategy, using Moore swabs might be sufficient if the goal is integrated surveillance for a broad set of bacterial, viral, protozoan, and helminthic pathogens. Improved environmental surveillance strategies can prepare and inform necessary health responses to infectious disease outbreaks (40) especially in LMICs where health care resources are limited and treatment is often not sought. The methods presented here provide a path towards a scalable integrated enteric pathogen surveillance strategy in communities in LMICs that lack networked sanitation infrastructure.

## Acknowledgments

This work was funded by the National Institutes of Health award R01AI155739.

We give our heartfelt thanks to the population of the commune of Comé for their enthusiastic participation in our studies, and in particular the Heads of school, the Households and the local health and administrative authorities who agreed to take part in the study. We are also extremely grateful to the IRD researcher collaborators for their confidence and for their continuous and generous support of our research efforts in Benin. We gratefully acknowledge the dedication and motivation of all members of the EPISOL study survey team, namely Félicien Chabi, Innocent Togbévi, Fidèle Houngbégnon, Fadel Tanimomon, Eloïc Atindégla and Elisée Adimi. We would like to thank the participants in the community from Timiri and Jawadhu Hills for their cooperation. We would also like to thank the district and state authorities for support and the permissions to carry out the study. We gratefully acknowledge the dedicated work carried out by all the EPISOL field workers and field and data managers - Mr. Rajeshkumar Rajendiran, Mr. Chinnaduraipandiyan Paulsamy and Mr. Gideon John Israel. We are very thankful to the WTRL biorepository for processing and banking these samples – Mr. Dhasthagir Basha and Mr. Sundarapandi.

## Supplemental

**S1 Fig. Positive and negative control qPCR results.** qPCR results for positive and negative controls. Internal Amplification Controls (IAC) - (pDMD801 in India or *B. atrophaeus* and *A. lumbricoides* known positive samples in Benin) underwent the DNA extraction protocol. Other control qPCR results include negative, no template controls (NTC), no samples extraction control (NSC), and positive standard controls. Standards were species-specific plasmids (10 pg, 100 fg, and 1 fg) in India or a single plasmid with all species-specific targets (1 fg) in Benin. Samples with failed internal amplification controls (IACs) were removed from our analysis.

**S2 Fig. Species-specific breakdown of positive qPCR control results.** qPCR results for species-specific standard controls. Standards were species-specific plasmids (10 pg, 100 fg, and 1 fg) in India or a single plasmid with all species-specific targets (1 fg) in Benin.

**S3 Fig. Field blank control qPCR results.** Species-specific qPCR results for wastewater blanks obtained from pouring bottled water into a whirlpack and then following extraction and qPCR protocol as described for grab samples. Internal amplification controls (IAC) (pDMD801 or *B. atrophaeus)* were spiked into samples during DNA extraction.

**S1 Table. Primers and probe sequences for qPCR**

**S2 table. Cycling conditions for TaqMan array card assays**

**S3 Table. Pathogen targets included in TaqMan array card**

