## Supplemental Information for "Environmental surveillance of soil-transmitted helminths and other enteric pathogens in settings without networked wastewater infrastructure"

*equal contributors as co-first authors

S1 Table. Primers and probe sequences for qPCR

| Species | Primers & Probes | Sequence (5’ - 3’) | Reference | Country |
| --- | --- | --- | --- | --- |
|  | Final concentration |  |  |  |
| *Ascaris lumbricoides* | Fwd  62.5 nM | 5’-CTTGTACCACGATAAAGGGCAT-3’ | (1) | Benin  India |
|  | Rev  62.5 nM | 5’-TCCCTTCCAATTGATCATCGAATAA-3’ |  |  |
|  | Probe  125 nM | 5’-/5-YakYel/TCTGTGCAT/ZEN/TATTGCTGCAATTGGGA/3IABkFQ/-3' | (1) Changed FAM to Yakima Yellow for Multiplex |  |
| *Trichuris trichiura* | Fwd  62.5 nM | 5’- GGCGTAGAGGAGCGATTT -3’ | (2) | Benin  India |
|  | Rev  250 nM | 5’- TACTACCCATCACACATTAGCC -3’ |  |  |
|  | Probe  125 nM | 5'-/5YakYel/TTTGCGGGC/ZEN/GAGAACGGAAATATT/3IABkFQ/-3 | (2) Changed FAM to Yakima Yellow for Multiplex |  |
| *Ancylostoma duodenale* | Fwd  500 nM | 5'-GTATTTCACTCATATGATCGAGTGTTC-3’ | (2) | Benin |
|  | Rev  500 nM | 5’- GTTTGAATTTGAGGTATTTCGACCA -3’ |  |  |
|  | Probe  125 nM | 5'-/56-FAM/TGACAGTGT/ZEN/GTCATACTGTGGAAA/3IABkFQ/-3' |  |  |
| *Necator americanus* | Fwd  250 nM | 5’-CCAGAATCGCCACAAATTGTAT-3’ | (2) | Benin  India |
|  | Rev  250 nM | 5’-GGGTTTGAGGCTTATCATAAAGAA-3’ |  |  |
|  | Probe  125 nM | 5'-/56-FAM/CCCGATTTG/ZEN/AGCTGAATTGTCAAA/3IABkFQ/-3' |  |  |
| *Bacillus atrophaeus* | Fwd  250 nM | 5'-GTCGTGACGCCAAATCTTCTC -3' | ZeptoMetrix LLC Catalog # 0801824 | Benin |
|  | Rev  250 nM | 5'-GGCTGTCAGACGATTATGCATTGA-3' |  |  |
|  | Probe  125 nM | 5'-/ABY/ATGCCGCCTTTTCCTCTT-3' |  |  |
| Internal Amplification Control (IAC) - pDMD801 | Fwd  250 nM | 5’-CTAACCTTCGTGATGAGCAATCG-3’ | (3) | India |
|  | Rev  250 nM | 5’-GATCAGCTACGTGAGGTCCTAC-3’ |  |  |
|  | Probe  125 nM | 5’- /56-FAM/ AGCTAGTCG/ZEN/ATGCACTCCAGTCCTCCT/3IABkFQ/ -3’ |  |  |

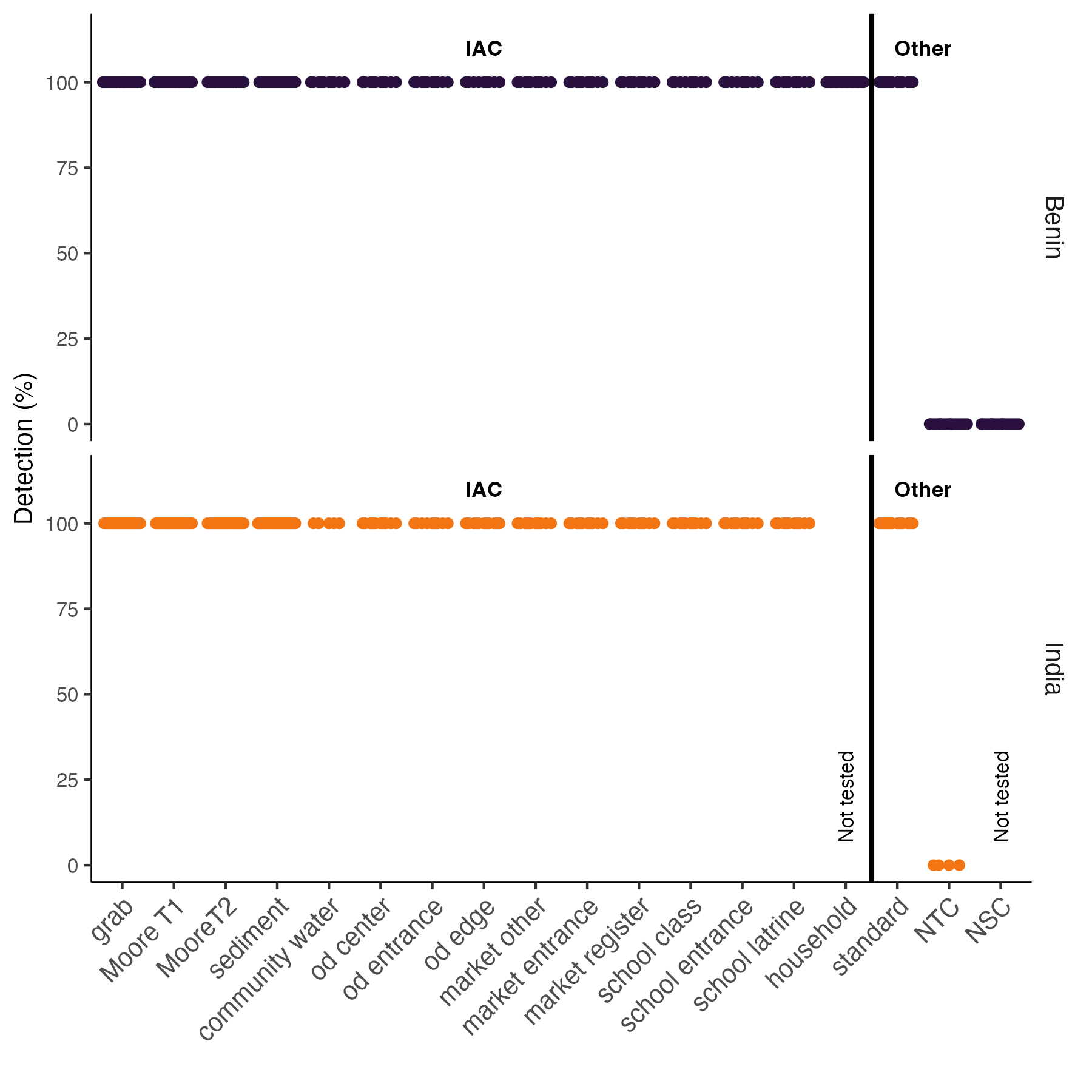

S1 Fig. Positive and negative control qPCR results. qPCR results for positive and negative controls. Internal Amplification Controls (IAC) - (pDMD801 in India or *B. atrophaeus* and *A. lumbricoides* known positive samples in Benin*)* underwent the DNA extraction protocol. Other control qPCR results include negative, no template controls (NTC), no samples extraction control (NSC), and positive standard controls. Standards were species-specific plasmids (10 pg, 100 fg, and 1 fg) in India or a single plasmid with all species-specific targets (1 fg) in Benin. Samples with failed internal amplification controls (IACs) were removed from our analysis.

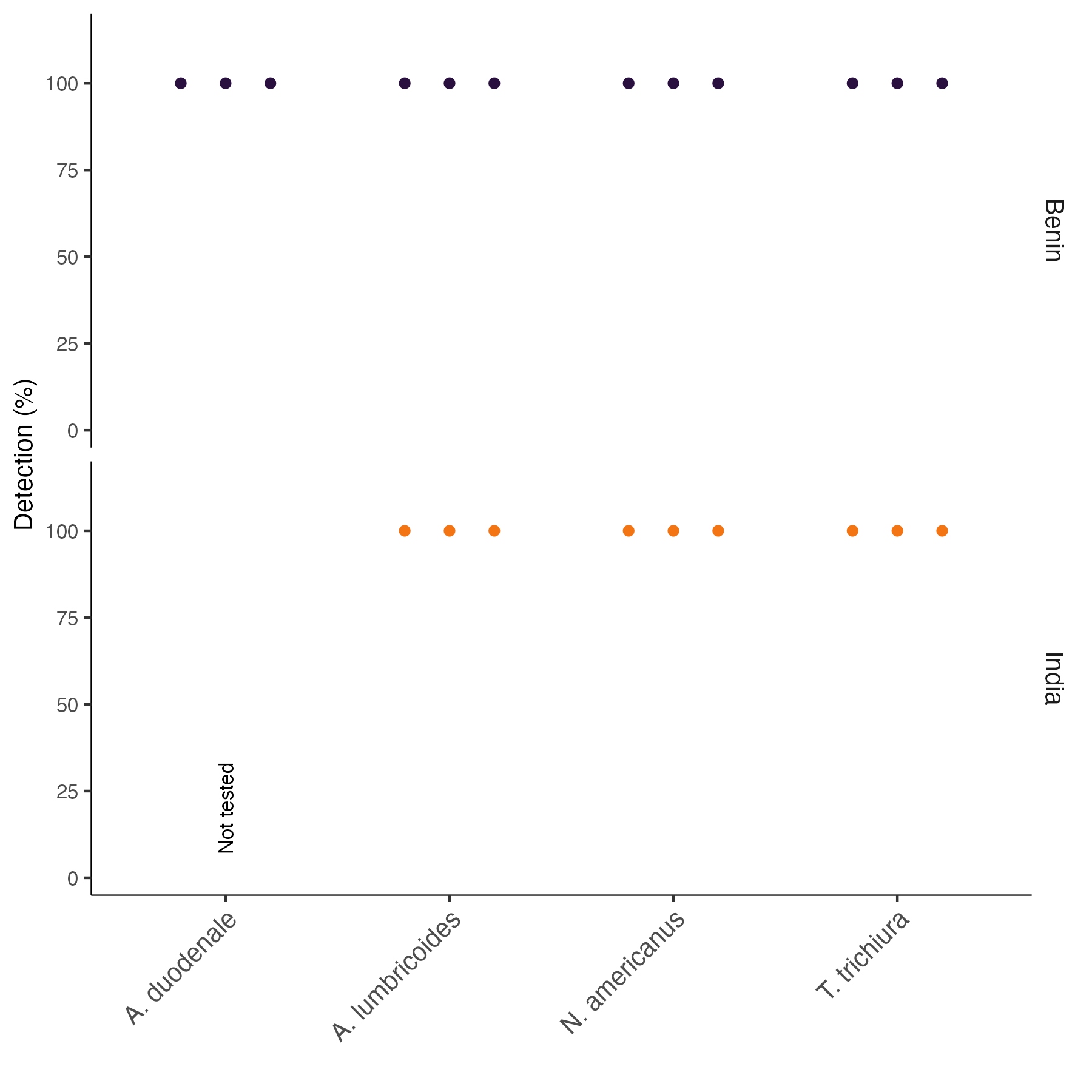

S2 Fig. Species-specific breakdown of positive qPCR control results. qPCR results for species-specific standard controls. Standards were species-specific plasmids (10 pg, 100 fg, and 1 fg) in India or a single plasmid with all species-specific targets (1 fg) in Benin.

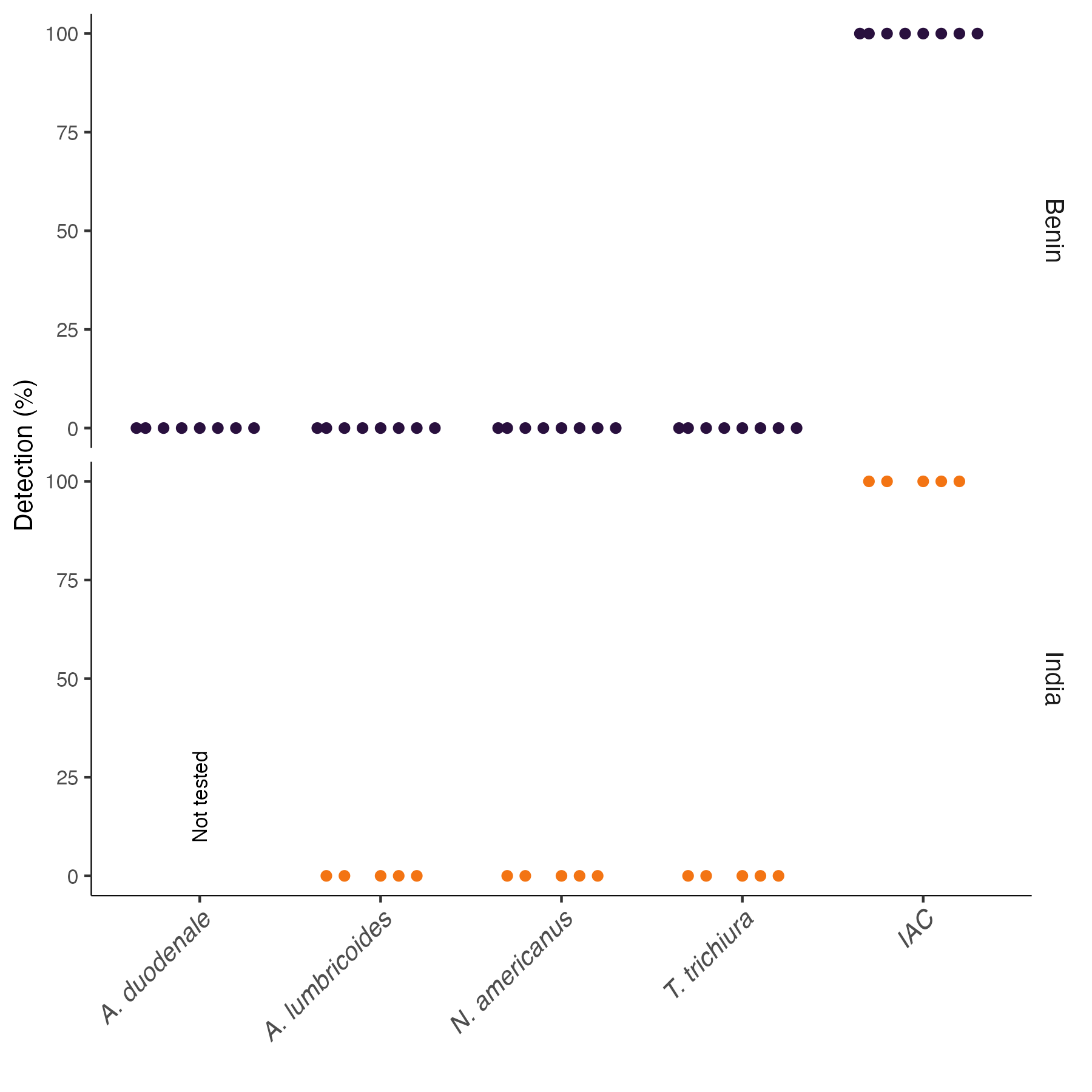

S3 Fig. Field blank control qPCR results. Species-specific qPCR results for wastewater blanks obtained from pouring bottled water into a whirlpack and then following extraction and qPCR protocol as described for grab samples. Internal amplification controls (IAC) (pDMD801 or *B. atrophaeus)* were spiked into samples during DNA extraction.

S2 table. Cycling conditions for TaqMan array card assays

| Hold stage | 95°c | 20 mins |
| --- | --- | --- |
|  | 95°c | 10 mins |
| PCR stage 40 cycles | 95°c | 15 secs |
|  | 60°c | 1 min |

S3 Table. Pathogen targets included in TaqMan array card

| Table 3. Pathogen and control targets for multiplex qPCR using custom TaqMan Array Cards. | | | |
| --- | --- | --- | --- |
| Category  (N targets) | Pathogen | Target (Benin) | Target (India)(4) |
| Helminth | *Ascaris lumbricoides* | Repetitive sequence(1) | *ITS1**(5)* |
|  | *Necator americanus* | Repetitive sequence(2) | *-* |
|  | *Trichuris trichiura* | Repetitive sequence(2) | *18S**(5)* |
|  | *Strongyloides spp.* | Repetitive sequence(6) | *-* |
| Protozoa | *Cryptosporidium spp.* | *18S**(5,7)* | *18S**(5)* |
|  | *Entamoeba histolytica* | *Ribosomal RNA**(8)* | *18S**(5)* |
|  | *Cyclospora* | *ITS1**(9)* | *-* |
|  | *Giardia duodenalis* | *GDH**(9)* | *18S**(5)* |
| Bacteria | ETEC | *Sth, Stp, LT**(10,11)* | *LT, ST**(5)* |
|  | STEC | *stx1, stx2**(11–13)* | *stx1 , stx2**(5)* |
|  | EPEC | *eae, bfpA* *(11,13)* | *eae, bfpA(5)* |
|  | EAEC | *aggR, aaiC, aatA(6,14,15)* | *aaiC, aatA(5)* |
|  | EHEC 0157:H7 | *rfbE(16)* | *rfbE(16)* |
|  | *Esherichia coli* | *uidA(17)* | *-* |
|  | *Escherichia coli/Shigella* | *ipaH(18)* | *ipaH(5)* |
|  | *Plesiomonas shigelloides* | *hugA(19)* | *-* |
|  | *Shigella flexneri* | *O-antigen**(20)* | *-* |
|  | *Salmonella typhi* | *Sty, tvib(21,22)* | *invA(5)* |
|  | *Helicobacter pylori* | *ureC(23)* | *ureC(23)* |
|  | *Campylobacter jejuni/coli* | *cadF, hipO, GlyA(24,25)* | *cadF(5)* |
|  | *Vibrio cholerae* | *toxR, ctxA(26)* | *-* |
|  | *Aeromonas* | *aha1**(27)* | *-* |
|  | *Bacteroides fragilis* | *HF183, HF134, BsteriF1**(28,29)* | *-* |
|  | *Clostridium difficile* | *tcdB(30)* | *-* |
|  | *Yersinia enterocolitica* | *virf(31)* | *-* |
| Virus | Adenovirus | *FIBER, hexon(6)* | *Hexon(5)* |
|  | Astrovirus | *capsid**(6)* | *-* |
|  | Enterovirus' | *5’ UTR**(6)* | *-* |
|  | Norovirus | *NGI**(6)* | *ORF1-ORF2**(5)* |
|  | Rotavirus | *NSP3**(6)* | *NSP3**(5)* |
|  | Sapovirus | *-* | *RdRp(5)* |
| Controls | 16S (total bacteria) | *16S**(32)* | 16S(5) |
|  | 18S | *18S* | *18S* |
|  | CrAssphage | *orf00024**(33)* | - |
